# Tackling the challenges of group network inference from intracranial EEG data

**DOI:** 10.1101/2022.10.25.513590

**Authors:** Anna Pidnebesna, Pavel Sanda, Adam Kalina, Jiri Hammer, Petr Marusic, Kamil Vlcek, Jaroslav Hlinka

## Abstract

Intracranial EEG (iEEG) data is a powerful way to map brain function, characterized by high temporal and spatial resolution, allowing the study of interactions among neuronal populations that orchestrate cognitive processing. However, the statistical inference and analysis of brain networks using iEEG data faces many challenges related to its sparse brain coverage, and its inhomogeneity across patients. We review these challenges and develop a methodological pipeline for estimation of network structure not obtainable from any single patient, illustrated on the inference of the interaction among visual streams using a dataset of 27 human iEEG recordings from a visual experiment employing visual scene stimuli. 100 ms sliding window and multiple band-pass filtered signals are used to provide temporal and spectral resolution. For the connectivity analysis we showcase two connectivity measures reflecting different types of interaction between regions of interest (ROI): Phase Locking Value as a symmetric measure of synchrony, and Directed Transfer Function - asymmetric measure describing causal interaction. For each two channels, initial uncorrected significance testing at p<0.05 for every time-frequency point is carried out by comparison of the data-derived connectivity to a baseline surrogate-based null distribution, providing a binary time-frequency connectivity map. For each ROI pair, a connectivity density map is obtained by averaging across all pairs of channels spanning them, effectively agglomerating data across relevant channels and subjects. Finally, the difference of the mean map value after and before the stimulation is compared to the same statistic in surrogate data to assess link significance.

The analysis confirmed the function of the parieto-medial temporal pathway, mediating visuospatial information between dorsal and ventral visual streams during visual scene analysis. Moreover, we observed the anterior hippocampal connectivity with more posterior areas in the medial temporal lobe, and found the reciprocal information flow between early processing areas and medial place area. To summarize, we developed an approach for estimating network connectivity, dealing with the challenge of sparse individual coverage of intracranial EEG electrodes. Its application provided new insights into the interaction between the dorsal and ventral visual streams, one of the iconic dualities in human cognition.

## 1 INTRODUCTION

Brain network dynamics give rise to the unique ability of (human) brain to carry out complex processing of external inputs and control complex behavioral outputs. To better understand the structure and dynamics of these networks, the tools of network neuroscience (Bullmore and Sporns, 2009) are being extensively developed and applied, ranging from efficient data acquisition and preprocessing to estimation and further analysis of brain connectivity networks. Multiple methods are available for neuroscientists to gain various types of insight into brain connectivity. Structural imaging, in particular diffusion weighted magnetic resonance imaging allows the estimation of the *structural connectivity* of the human brain using the methods of tractography, while various methods of functional brain imaging, including in particular the functional magnetic resonance imaging (fMRI) and electroencephalography (EEG), enable the estimation of the patterns of statistical dependencies between remote neurophysiological events, known as the *functional connectivity* (Friston et al., 1993). While the functional connectivity provides in principle only information that the activity of some brain areas is related, a whole plethora of advanced methods for the estimation of the directed, cause-effect relationships between brain areas has been developed, commonly summarized under the term *effective connectivity* (Friston, 1994).

Inferring the effective connectivity can be seen as providing the ultimate answer to the question of information flows in the brain, however effective connectivity estimation is hindered by a range of technical challenges. Successfully tackling these challenges would provide an appropriate causal description of the brain network, that would lend itself to further processing using the tools of complex network theory.

In this work, we discuss a range of such challenges and possible solutions, presenting an analysis of a particularly problematic dataset using a pipeline implementing a specific combination of solutions. Note that a combinatorial explosion of the number of possible pipelines makes a comparison of all alternatives not feasible - thus the presentation serves as an example rather than a comparison of all tools available. The challenging task that we select for the demonstration is the estimation of causal network from intracranial EEG (iEEG). Intracranial EEG recording as such is a powerful tool for the discovery of brain function. This type of measurement is characterized by both high temporal and spatial resolution, a combination which is impossible for other, non-invasive, neuroimaging techniques. However, the analysis of the iEEG data faces a set of challenges mainly because of in-homogeneous and spatially sparse data acquisition - the measurement sites, corresponding to the implanted electrodes, are few and far between, thus not covering all the areas of interest in any one particular subject. Moreover, the positioning of the implantation differs widely between subjects, thus complicating a group-level analysis. In our paper, we develop the methodological pipeline for estimation of such group network structure, that can’t be estimated from any single patient. As an example, we show the connectivity analysis of the visual processing streams.

### 1.1 Visual streams conundrum

Large body of evidence suggests that visual information is processed in the brain in two separate streams originating from the visual cortex: the dorsal and ventral streams in parietal and temporal lobes respectively. These streams were originally characterized in terms of the “where” and “what” distinction (Ungerleider and Mishkin, 1982), suggesting the role of the dorsal stream in spatial position and motion coding and of the ventral stream in object identity coding. Examination of patients suffering visual agnosia after bilateral ventral lesions, like D.F., in combination with their unimpaired spatial-motoric abilities lead to an alternative model (Goodale and Milner, 1992). In this model, characterized as “how” and “what” model, the dorsal and ventral streams are distinguished not by the type of information processed, but by the results of their visual processing, which is motoric action or conscious perception. According to this view, the dorsal stream processes the visual information to enable movements in space like grasping an object, while the ventral stream processes the same information for perception of objects, their identity and also their spatial relationship. Several brain areas in the ventral stream are specialized for recognition of various classes of visual percepts. For instance, damage to an area in lingual gyrus leads to landmark agnosia (Aguirre and D’Esposito, 1999) and a more anterior parahippocampal place area (PPA) in the posterior parahippocampal/anterior lingual region is one of the brain regions specifically responding to spatial scenes (Epstein and Kanwisher, 1998).

In contrast to the well established role of ventral stream, there are still controversies regarding the dorsal stream function and its connections with the ventral stream. The dorsal stream actually seems to divide into three parallel pathways serving different functions. The most ventral of them, the parieto-medial temporal pathway, has been associated with spatial information transfer from dorsal to ventral stream. The existence of this pathway was suggested based on monkey and human brain anatomy (Kravitz et al., 2011) and is also supported by human resting-state functional connectivity (Boccia et al., 2017). Its functional properties have been suggested based on the role of the included structures like retrosplenial cortex (Byrne et al., 2007). Besides this pathway, the dorsal and ventral streams seem to be interconnected several times, with the dorsal stream providing attention focus and spatial information (Cloutman, 2013). On the other hand, the occipito-temporal cortex in the ventral stream seems to provide object identity information to the dorsal stream as was only indirectly suggested from a human fMRI experiment (Kristensen et al., 2016). Direct evidence about these connections and their functional role is however still scarce.

## 2 CAUSAL NETWORK INFERENCE

Many challenges of causal network inference are generic and relevant irrespective of the method used to obtain the time series. For instance, in principle, any estimation method is negatively affected by the presence of measurement noise and estimation from only short observations, issues present to various extent in all neuroimaging methods. Indeed, standard methods such as the common linear implementation of Granger causal analysis, that rely on assumptions concerning the underlying process (namely, that it is a stationary linear autoregressive process), may provide relative advantage in sensitivity for short time series. This advantage against nonlinear methods with less or no assumptions may be practically relevant even for systems that, in principle, have nonlinear character – such as the brain or the Earth’s climate (Hlinka et al., 2013).

However, the character of neuroimaging data poses additional challenges beyond its nonlinearity, ranging from some very general to those quite specific to iEEG data. A first commonly faced challenge is the multivariate, rather than bivariate, character of the data. Indeed, in principle one can estimate the causal interactions separately for each pair of variables. The downside of such straightforward approach is the computational demands, but more importantly, the fact that the estimated interaction between any pair of variables might be confounded by not accounting for the effect of other variables in the role of mediators or common drivers. If one believes that the other variables observed may play such a role, it is advantageous to fit a joint multivariate model that takes (all) these variables into account. Of course, such a model has a large number of parameters, and might thus call for even longer time series samples, and pose additional computational problems, particularly when nonparametric methods, such as conditional mutual information, are in use. Then, an alternative approach lies in the stepwise inference of a smaller set of causally relevant ‘parent’ variables for each target variables - see Kořenek and Hlinka (2020) for a systematic comparison of a set of such state-of-art methods.

A second challenge lies in the nonlinear character of the brain dynamics, which calls for the application of nonlinear, potential nonparametric methods, such as the conditional mutual information. However, a range of prior studies have shown, that the nonlinear/nongaussian character of neuroimaging data may be relatively weak, and thus the use of linear methods might be justified in terms of the trade-off between the lost information and higher speed and general reliability of linear approaches - see a discussion for fMRI (Hlinka et al., 2011; Hartman et al., 2011) and electroencephalography (Blinowska and Malinowski, 1991). Therefore, the choice of linear methods is generally suitable.

A third aspect that must be considered with brain signals, particularly with EEG, is that the dynamics and interactions take place across a range of time scales. In fact, specific EEG activity frequencies have been shown to be related to specific types of cognitive or arousal states. Therefore the use of frequency-specific estimates of causal interactions is common. The downside of this approach is that the inferred object further increases dimension - causality is resolved not only across all pairs of observed signal channels, but also across frequencies. We shall discuss the consequences of this aspect for overall statistical inference later.

Albeit the multivariate character of the observed data allows (and calls for) taking into account indirect causality mediated/driven by any of the observed variables, the effect of any other, directly unobserved, variables is not accounted for in our methodology, and this omission may lead to inference of spurious causal links. It is important to highlight that this is a generic effect that affects most of the available methods for causal inference. While some methods have been proposed to suppress the effect of such unobserved (latent) variables (such as those based on the utilization of time reversal (Haufe et al., 2012; Winkler et al., 2016)), their assumptions are not universally fulfilled, and particularly in multivariate setting can lead to wrong conclusions (Kořenek and Hlinka, 2021). Although other methods are available that attempt to suppress the effect of latent variables and non-neuronal interactions, such as the volume conduction in tissue, the problem is far from ultimately solved, and one should always interpret the results of causal analysis with caution in terms of the potential role of any latent intervening variables.

Another complication, which is not entirely specific, but quite common in EEG analysis, is that the systems is deemed to be substantially nonstationary, and thus the structure of causal interactions may meaningfully change in time. A typical engineering solution to such situation is to estimate the causal structure using a (potentially overlapping) sliding window for a sequence of time periods. Note that an alternative exists in using variants of Kalman filtering (Kalman, 1960) - this avoids the problem of selecting the window parameters; however still requires selection of other intrinsic parameters.

The requirement of obtaining time resolved causality leads to estimation from short time series; to compensate for the loss of sample size, in the case of task-based iEEG, one may typically utilize data across many trials for the estimation of the parameters of the underlying vector autoregressive process. The consequence of this is that only a single estimated causal structure is estimated across all trials. The trial dimension can not be thus used in straightforward way further statistical assessment of the inferred causal links, i.e. by evaluating the distribution across trials with respect to the posited null hypotheses. However, alternative ways for establishing empirical null distribution are available, as discussed below, that are based on the use of bootstrapping, surrogate data or permutation testing.

Indeed, the evaluation of the statistical significance of causal analysis results obtained by most methods of causal inference relies on some construction of empirical null distribution of selected test statistics. The distribution of the causal indices under the hypothesis of no interaction is, however, in most cases not known analytically, see e.g. the approach of surrogate testing proposed in Kaminski et al. (2001).

Note that with task-based electrophysiology data, we might not necessarily focus on the structure of causal interactions per se, but on the deviation of the structure from the resting baseline. The rationale is two-fold. First, it can be reasonably expected that in the resting state, most parts of brain are functionally connected and the change in the task scenario might contribute only (potentially minor) alteration to this ongoing process of interactions (Biswal et al., 2010). Second, due to the latent variables as well as technical confounds and volume conduction, the causal structure estimated in task is expected to be biased towards detecting a range of spurious interactions. However, the same set of spurious sources are to affect the estimate of interactions outside the task condition of interest. Therefore, some form of correction for these effects by subtraction or comparison with respect to the baseline estimate of causal structure may be warranted to identify the causal interactions specific to the task condition of interest.

A specific problem mentioned earlier is the high dimensionality of the estimated object. In particular, the causal interaction is estimated between any two channels (taking all other channels into account), for each frequency and time window. Even with a valid statistical significance test, such situation invites a multitude of false positive results due to the multiple testing problem involved. Indeed, one may attempt to apply one of the methods for multiple testing correction, that controls the family-wise error rate (FWE), or, as a less strict option, the false discovery rate (FDR) (see Hochberg (1988); Benjamini and Hochberg (1995)). However, the application of these methods may decrease the overall statistical power, lead to false negatives and also create extreme computational demand, as p-value estimates with high precision call for the generation of high number of permutations/surrogates to generate the null distributions. A potential solution to this problem lies in some dimension reduction of the test statistics prior to its statistical evaluation.

Finally, there are challenges that are highly specific to the iEEG data. In particular, each subject has their own set of implanted electrode locations at different brain positions, that is moreover extremely sparse and inhomogeneous. How can one in such a situation reliably estimate the structure of brain network interactions across the whole group of subjects? A reasonable solution seems to be to (at least temporarily) sacrifice some of the spatial resolution (that is anyway somehow artificial with only sparse coverage), and aggregate the data within pre-defined generic brain regions. Note that the data can, with advantage, be aggregated (by averaging or with another approach) also later in the processing, such as at the level of connections, rather than already at the level of the measured signals.

Note that for task activation data, such as evoked potentials, an alternative approach would be to merge all the channels into one joint dataset, to obtain a single ‘super-subject’, while ignoring that the channels actually come from different subjects. However, in case of analysis of connectivity this approach is not directly applicable, because the analysis of connectivity between channels from different subjects might be detrimentally affected by the lack of synchronization of processes across subjects. Also, for some purposes, the interpretation of the brain network calls for some coarse-graining to the level of reproducible and interpretable brain regions, and thus rendering detailed info on individual channels not necessary.

The collating the brain connections into generic region pairs can in principle entail both integrating the data within subjects, as well as across subjects. Given the heterogeneous sampling over space and subjects, different regions pairs will have contributions from different number of channels across different subjects. Together with additional problem of dependence between channels (and channels pairs) within subjects, this poses another problem of statistical inference concerning the connections observed at the inter-regional level. We believe that the solution to this challenge lies in the construction of a surrogate data that respects the within-subject channel-pair dependence as well the contribution of particular subjects to the particular region connections. Note that an alternative solution has also been applied in the literature, that aimed for analysis of only those pairs of regions, that are sampled (at least by a single pair of channels) in some minimal number of subjects, that allows statistical inference across subjects (of some subject-level test statistics, typically obtained by summing/averaging within subject) (see, for example, Burke et al. (2013, 2014); Bastin et al. (2017)).

Below, we develop a pipeline for the analysis of the interaction of the two cortical visual processing streams, applied to a recorded iEEG task-based dataset used by Vlcek et al. (2020). In this analysis, we illustrate the considerations outlined above, that need to be taken into account while analysing the causal structure of such complex neuroimaging datasets.

## 3 DATA

The current study used a dataset including 27 patients, who voluntarily participated in a visual recognition experiment, as described by Vlcek et al. (2020). All patients had normal or corrected to normal vision. This study was approved by the Ethics Committee of Motol University Hospital and all patients gave their informed consent to participate. Intracranial EEG was recorded in the Motol Epilepsy Center in Prague in pharmacoresistant epilepsy patients using depth platinum electrodes (Dixi Medical), designed for local field potential (LFP) recordings. Electrodes were 0.8 mm in diameter, with 5-18 contacts of 2 mm length, separated by insulator segments of 1.5 mm length. Up to 128 contacts were recorded based on the medical requirements. Wideband signals (0.05-512 Hz) were recorded with a sample rate of 512, 2048, or 8000 Hz, and later subsampled to 512 Hz. All contacts were localized in the standard Montreal Neurological Institute (MNI) space.

Because the implantation is decided by clinical requirements, it naturally includes many areas not involved in visual processing. Thus, to limit computational costs and improve statistical power, only those channels with significant responses to demonstrated pictures were analyzed (see Vlcek et al. (2020) for statistical analysis details). Moreover, for the purposes of connectivity analysis, only those subjects, who had implanted at least two of the initially defined seven regions of interest (ROIs) relevant for the cognitive tasks studied (described in Sec. 4), were included in the analysis. This provided a subset of 15 subjects, the data of which were included in the final analysis (8 women, median age 30 years, range 17–48 years, education level: 1 primary school, 11 secondary school, and 3 college-level education).

Each time series was normalized by the baseline standard deviation. Bipolar montage was used to remove the influence of the common source, i.e. to ensure that the processed signal had a local origin. We refer to the bipolar contact pairs further as “channels”. To eliminate the non-stationarity of the time series, we computed the differences in time, i.e. from the signal *y_t_* we computed 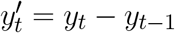 (Barnett and Seth, 2014; Seth, 2010).

### 3.1 Visual experiment

During the visual recognition experiment, four categories of pictures were demonstrated to the participants on a computer screen: scenes, small objects of daily life, faces, and finally a control category of edibles (fruits or vegetables). Images of the last category were used to control for a potential attention decrease – subjects were instructed to press a button every time they see a picture of this category.

Each of the categories of interest (scenes, objects, faces) included 100 different pictures. Each of these was shown in two different trials, giving the total of 200 trials per category, in pseudo-random order. The number of different pictures of edibles (fruits/vegetables) was 25, each being shown twice giving rise to 50 trials in total. Stimuli were presented for 300ms, each followed by 800ms of a black screen with a white cross in the centre. For the analysis, the data were cut so that every trial had 200ms of baseline (the black screen before the stimulus demonstration), 300ms of picture demonstration, and 500ms of the following black screen. Trials with fruits/vegetables, as well as with button presses by mistake were excluded from the analysis. Also, trials with epileptogenic activity were excluded. A detailed description of stimuli, the task and the procedure used for removing the data with epileptogenic activity can be found in Vlcek et al. (2020).

### 3.2 ROI definitions

A previous study on the same dataset (Vlcek et al., 2020) has defined 7 clusters of channels that were of particular interest as they have shown a stronger reaction to the visual stimulation by scenes than by objects. These thus define effective regions of interest (ROIs), the connectivity among which we shall attempt to establish.

However, connectivity analysis on that subset of channels might be too restrictive because of the limited coverage of the visual pathways and consequently a relatively small amount of edges that could be tested. Therefore, we extended the dataset with all the channels that reacted to any type of stimuli, and lied close to the previously defined clusters. In particular, we constructed a 3D convex hull of every cluster; and extended every hull to 0.5-4.5 mm millimetres such that corresponding regions have more uniform volumes. Those increased convex hulls then defined, which channel belong to which ROI. Number of implanted channels per ROI can be found in Table 1. Note that we didn’t distinguish the hemispheres, i.e., we consider every ROI symmetric, laying in both hemispheres.

**Table 1.** Number of implanted channels per ROI for analyzed subjects.

| Subject | ROI 1 | ROI 2 | ROI 3 | ROI 4 | ROI 5 | ROI 6 | ROI 7 | Total |
| --- | --- | --- | --- | --- | --- | --- | --- | --- |
| Subject 1 | - | - | 2 | 1 | - | 3 | - | 6 |
| Subject 2 | - | - | 2 | 1 | - | - | - | 3 |
| Subject 3 | - | - | - | 2 | - | - | 6 | 8 |
| Subject 4 | - | 1 | 1 | - | - | 2 | - | 4 |
| Subject 5 | - | - | 7 | 1 | - | - | 1 | 9 |
| Subject 6 | - | 5 | - | - | - | - | 1 | 6 |
| Subject 7 | 2 | 2 | - | - | - | 4 | - | 8 |
| Subject 8 | - | - | - | 4 | - | - | 4 | 8 |
| Subject 9 | - | - | - | 3 | - | - | 1 | 4 |
| Subject 10 | 1 | - | - | - | 1 | - | - | 2 |
| Subject 11 | - | - | 1 | 3 | - | 1 | - | 5 |
| Subject 12 | 6 | - | - | - | 6 | - | - | 12 |
| Subject 13 | - | - | 3 | 2 | - | - | 3 | 8 |
| Subject 14 | 4 | - | - | - | 3 | - | - | 7 |
| Subject 15 | - | - | 2 | - | - | - | 1 | 3 |
| **Total** |  | **8** | **18** | **17** | **10** | **10** | **17** | **93** |

The centroids of these clusters were localized in the following structures:

1. OPA - the posterior angular and middle occipital gyrus (MNI [38, −76, 24], Occipital Place Area),
2. pLG - the posterior collateral sulcus at the junction with the lingual sulcus (MNI [25, −72, −8]),
3. PPA - the lingual and fusiform gyrus along the middle collateral sulcus (MNI [30, −45, −7]], Parahippocampal Place Area),
4. aCOS - the parahippocampal and fusiform gyrus along the anterior collateral sulcus (MNI [32, −26, −22]),
5. PCUN - the precuneus (MNI [15, −61, 27],
6. MPA - the superior part of the lingual gyrus and precuneus next to the retrosplenial region (MNI [16, −53, 12], Medial Place Area),
7. HIP - the anterior hippocampus (MNI [25, −12, −20]).

## 4 HYPOTHESIS

For the purpose of our current analysis, we developed a hypothesis about the connectivity between seven ROIs (Figure 2), based on our previous iEEG study (Vlcek et al., 2020) and other electrophysiological and anatomical published results (Kravitz et al., 2011; Henriksson et al., 2019; Baldassano et al., 2016; Bastin et al., 2013; Boccia et al., 2017; Cavanna and Trimble, 2006; Epstein and Baker, 2019; Byrne et al., 2007).

**Figure 1.**
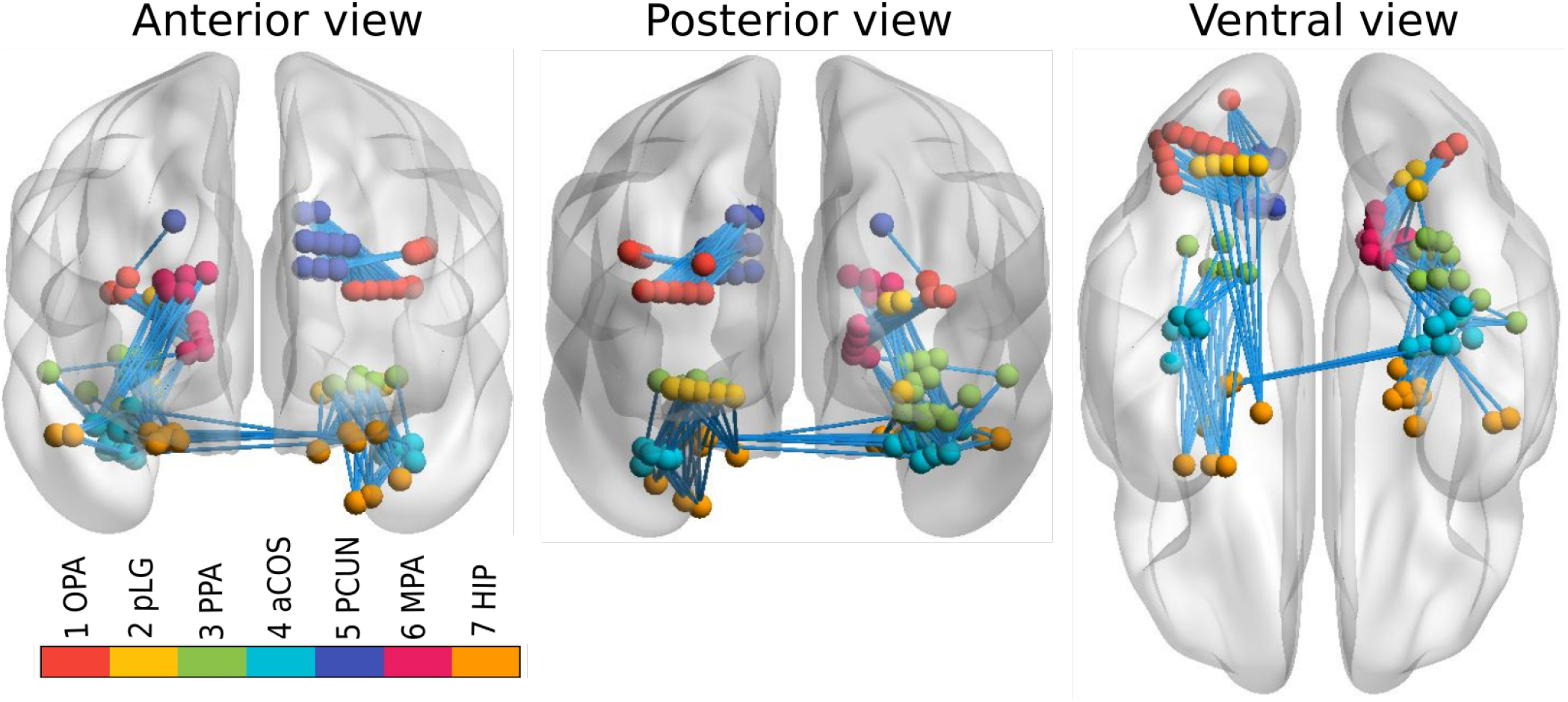
Implanted channels (aggregated across all subjects) with ROIs marked by different colours. Connections between ROIs, available for testing, are shown by light blue lines.

**Figure 2.**
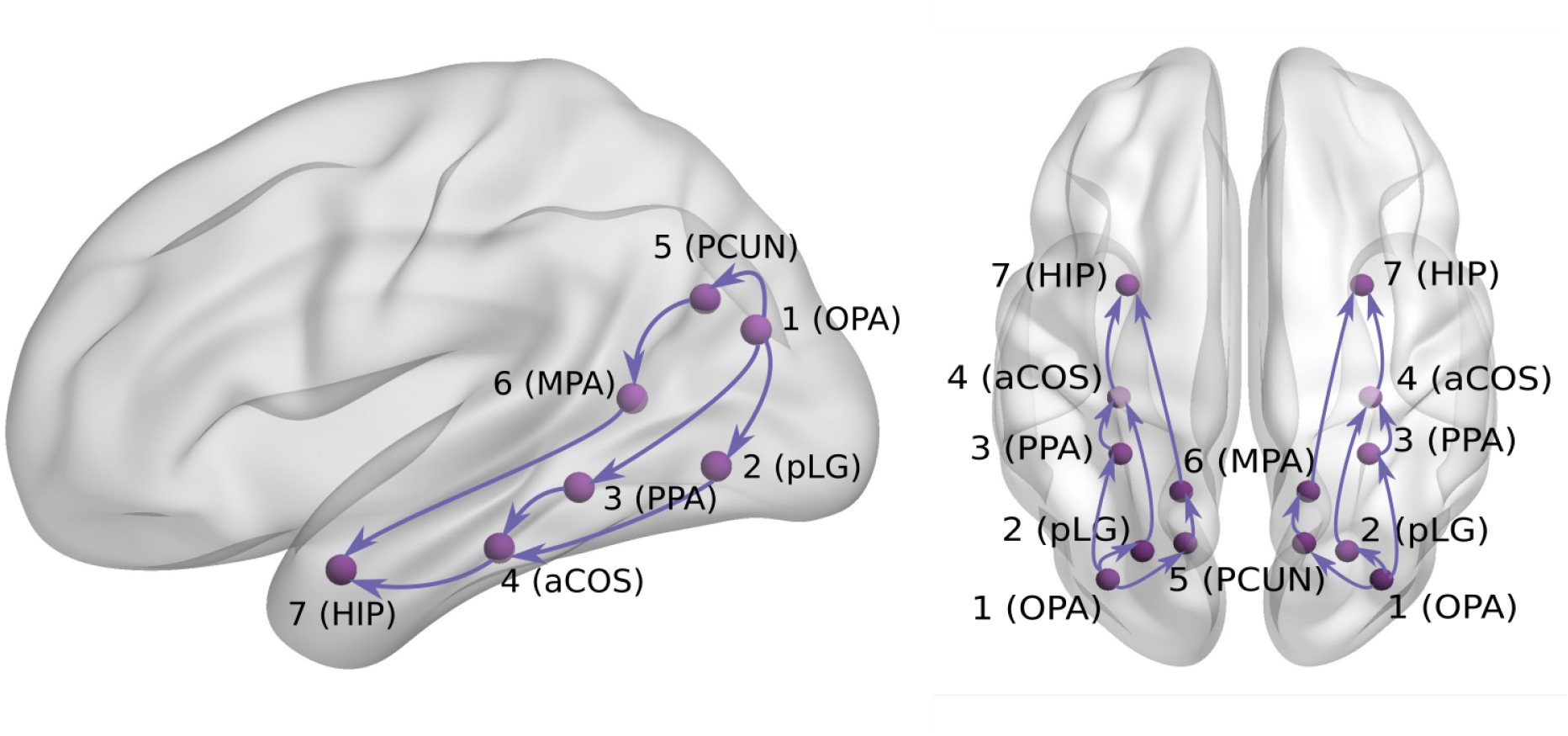
Hypothesis about visual signal spreading via ventral/dorsal visual pathways.

First, the visual cortex areas project to parieto-occipital areas such as the Occipital Place Area (OPA, ROI 1) around the transverse occipital sulcus (Kravitz et al., 2011). The documented responses as early as 60ms after scene presentation (Henriksson et al., 2019) support its position early in the visual scene analysis network. As our data sample did not contain any primary visual cortex channels, we considered the OPA to be the entry point of the studied network of 7 regions.

Second, the OPA is functionally connected with the Parahippocampal Place Area (PPA, ROI 3) in the posterior temporal cortex (mainly its posterior part (Baldassano et al., 2016)). Several studies documented an early scene selectivity in the PPA at around 150-200ms (Vlcek et al., 2020; Bastin et al., 2013). Based on the results of Vlcek et al. (2020), we distinguished another early scene selective area in the lingual gyrus (pLG, ROI 2). Because of its large selective responses to scenes but a lack of published information about its role in scene analysis, we located it in our hypothetical network in parallel to the PPA area. Therefore, both PPA (ROI 3) and pLG (ROI 2) are hypothesized to receive input from ROI 1, with unclear interaction between ROI 2 and 3.

Next, the resting state fMRI data suggest the PPA area connects with more anterior temporal regions including the anterior hippocampus (ROI 7) (Boccia et al., 2017). In our previous study (Vlcek et al., 2020), we documented late anterior hippocampal scene selectivity starting at around 300-350ms and even earlier responses at about 200-250ms around anterior collateral sulcus, which is positioned anatomically between the PPA and hippocampus. We added this region, located between ROI 3 and 7, as ROI 4.

Previously, we documented the scene selectivity also in the dorsal visual stream. In the posterior precuneus, the scene selectivity appeared around 200-250ms (Vlcek et al., 2020), which is in agreement with its role in visual imagery and spatial attention (Cavanna and Trimble, 2006). Due to its location in the parietal cortex, we added it to our hypothetical network as ROI 5, parallel to ROIs 2 and 3.

Finally, the ROI 6 in our hypothesis corresponds to the Medial Place Area (MPA, also known as Retrosplenial Complex), a region with well documented scene selectivity also from fMRI studies (along with OPA and PPA, see Epstein and Baker (2019) for review). We previously documented scene selectivity in this region starting at around 300-350ms, later than in the posterior precuneus, but similarly late to the hippocampal responses. As this area was documented to be part of a parieto-medial temporal pathway (Kravitz et al., 2011), translating visuo-spatial information between the dorsal and ventral streams (Byrne et al., 2007), we added it to our hypothetical network, potentially mediating the information flow between the ROI 5 in the dorsal stream and ROI 7 in the ventral stream.

## 5 METHODS

### 5.1 Connectivity analysis: general approach

For the connectivity analysis purposes, we chose two different connectivity measures. The Phase Locking Value (PLV, (Lachaux et al., 1999), see Section 5.2) is an example a undirected measure of synchrony between two time series, while the Directed Transfer Function (DTF, (Kaminski and Blinowska, 1991), see Section 5.3) is one of most commonly used directed connectivity measures, i.e. it estimates directed interactions between channels.

As discussed earlier, implantation according to the clinical needs implies limitations on the collected data. For instance, connectivity between two regions can be naturally computed only for those patients, who have electrodes in both regions. Note that in principle one may attempt assessing dependence between signals from different regions in two different subjects and estimate thus some shared, population-level, component of the functional or even effective connectivity, and ultimately of the information flows, it is not entirely straightforward what interpretation would be suitable for such connectivity measure in view of inter-subject variability and noise. While at least at the level of correlations, inter-subject dependencies (even inter-species correlation) has been used e.g. in natural viewing data to identify homological areas across between monkey and human (Mantini et al., 2012), due to the complicated interpretability, we leave this as a matter of future research and restrict ourselves to pairs of channels observed within any single subject.

Figure 3 shows the number of implanted pairs of ROIs and channels. Ideally, we would like to analyse the data with all 7 ROIs sampled in the same patient, but we don’t have such subjects - at maximum, 3 ROIs were measured together. Also, as shown in Figure 3, not all links between ROIs were measured. Note that this is a typical situation, rather than an exception, in case of human iEEG studies. Thus, one is limited to analysing only the observable connections between the regions by calculating connectivity per subject and then collected links per pair of ROIs to form the group-level network connectivity graph.

**Figure 3.**
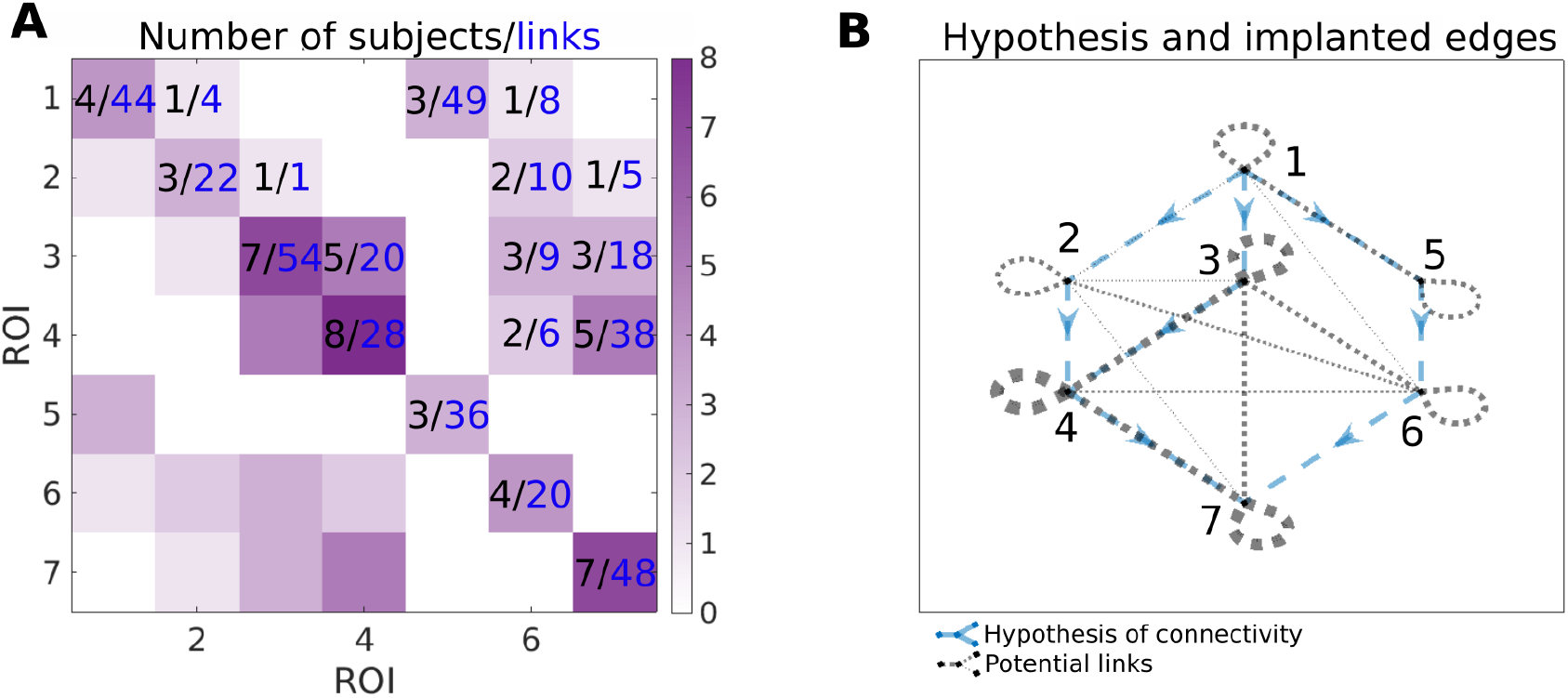
Information about the number of implanted patients (black numbers) and pairs of channels (blue numbers) for each pairs of ROIs. In the panel A, *ij*-th element of the matrix corresponds to the number of patients, that have implanted both ROI *i* and *j* (black value) and number of pairs of implanted channels spanning ROIs *i* and *j* (blue value). Color-scale corresponds to the number of patients with implanted pair of ROIs, i.e. it corresponds to the values in black. In the panel B, the graph of the hypotheses is combined with the graph of the connections that can be assessed from the data. The hypotheses are shown as blue dashed arrows. The implanted ROI pairs are connected with a black dotted edge, the width corresponds to the number of patients who have that pair of ROIs implanted. For example, for the connection from 3 to 4, the number of implanted subjects and pairs of channels can be found on the panel A (5 subjects, 20 pairs of channels); corresponding hypothesis and implanted edges can be found on the panel B.

The scheme of forming the group-level network connectivity graph from the subject-specific graphs is described in detail in Section 5.4. For a particular pair of channels (patient-specific), we compute the binary map of significance of the time- and frequency-resolved connectivity (see Figure 4). Significance testing for every time-frequency point is carried out using the multivariate Fourier surrogates (Prichard and Theiler, 1994), estimated from the baseline time series segment; the significance testing thus provides a binary time-frequency map of connectivity between the two channels.

**Figure 4.**
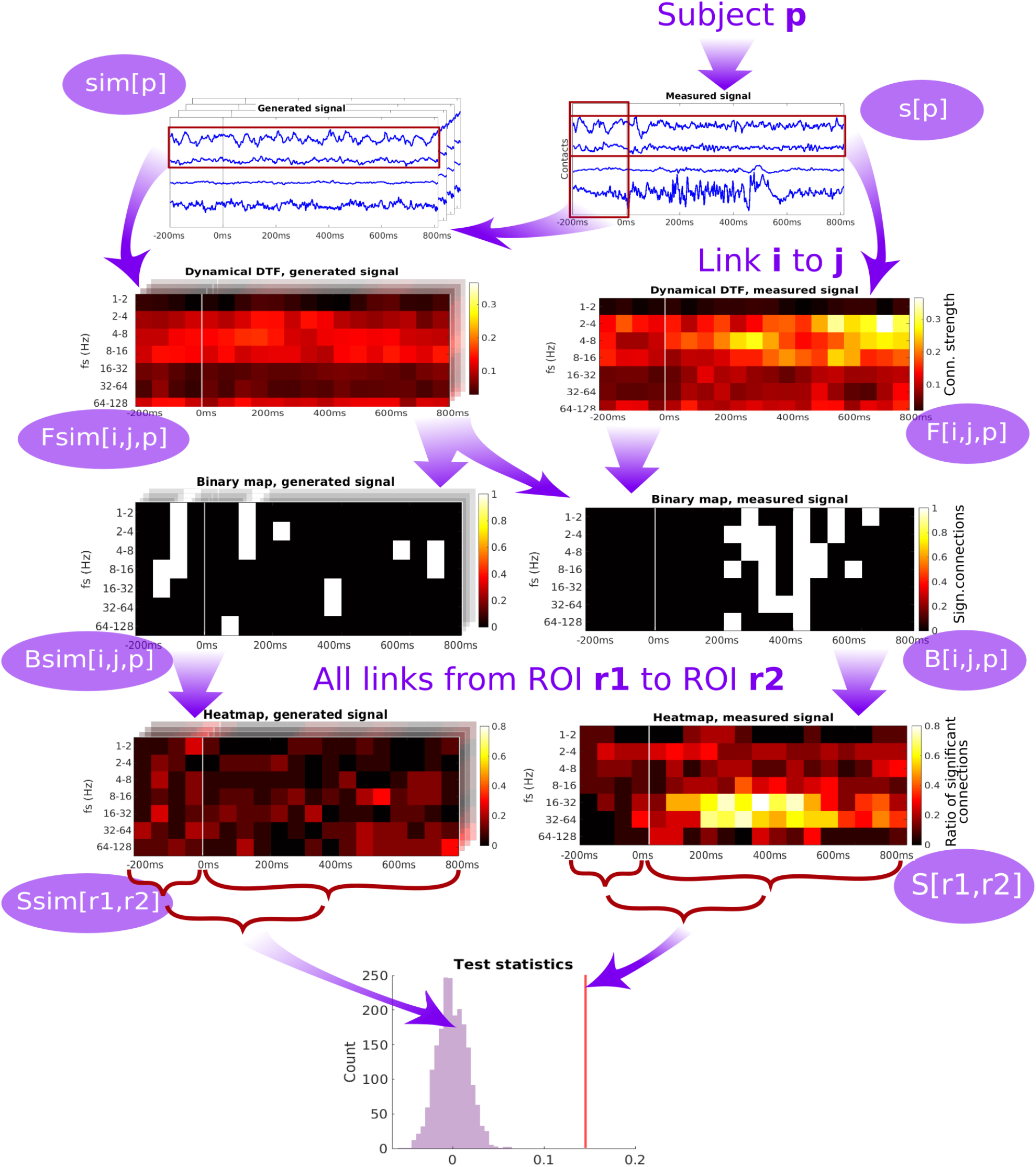
Diagram of the pipeline showing the subsequent steps starting from the time series measured from a single patient, through computation of dynamic connectivity, binary maps, heatmaps and test statistics. DTF is used as an example of connectivity measure.

Then, binary maps for all pairs of channels belonging to a given pair of ROIs are summed up (across all relevant channel pairs within a subject, and across all subjects) to form a ROI-pair specific connectivity density (*”heatmap”*, see Figure 4). While this heatmap gives a good qualitative overview of which connections are active at which time and frequency, carrying out multiple testing comparison across the many dimensions of the problem (time, frequency, channel pairs, subjects) might lead to severely decreasing the statistical power. To tackle this problem, as the connectivity can be expected to be relatively smooth over time and frequency (which is confirmed upon visual inspection), we chose to further proceed by conservatively evaluating the overall strength of the link between a pair of ROIs by computing an average heatmap value from the period after the stimulation.

The average heatmap thus captures the percentage of channel pairs, that showed significant connectivity at a given time and frequency bin. Notably, even in the case of no true connectivity, and in particular in the case of no difference from the baseline resting activity, one may expect some significant results. In theory, this percentage should correspond to the significance level selected for the channel-pair test. However, even though the standard approach of multivariate Fourier surrogates is used, the baseline data may violate its assumptions - particularly stationarity (and gaussianity), and thus having false positive rate increased above the nominal significance level even in the baseline segment. To further conservatively correct for this effect, we corrected the average heatmap value of the response period by subtracting the average heatmap value during the baseline. To provide a statistical inference regarding this final value of connectivity strength between ROIs, it is finally compared to a histograms of values of the same statistic obtained from multivariate surrogates of baseline data.

### 5.2 Connectivity measure: PLV

Phase Locking Value (PLV) was defined in Lachaux et al. (1999) and it is a time-dependent connectivity measure, that can be used to investigate task-induced (changes in) synchronization between time series. PLV is defined as

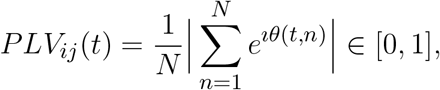

where *N* is a number of trials, *θ*(*t, n*) = *ϕ_i_*(*t, n*) – *ϕ_j_*(*t, n*), and *ϕ_i_*(*t, n*) and *ϕ_j_*(*t, n*) are the instantaneous phases for signals *s_i_* and *s_j_* in trial *n* at time *t*. In our study, the instantaneous phase of the signal was computed using a Hilbert transform. Higher values of PLV, with maximum of 1, mean higher synchronization between the time series (in terms of similar phase-lag between two signals across trials), while zero PLV corresponds to independent instantaneous phases. Before computing the PLV, the measured signal was filtered in several frequency bands, using a band-pass filter. Detailed pipeline is described in Section 5.4.

### 5.3 Connectivity measure: DTF

Directed Transfer Function (DTF) is a connectivity measure based on VAR(*p*) (Vector Autoregressive Process of the order *p*, see Lütkepohl (2005) for a general overview).

Let {*X_t_* ∈ ℝ^*n*^ : *t* = 1, …, *T*} denote a multivariate time series. The VAR(*p*) is defined by

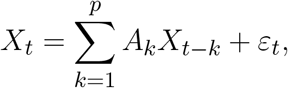

where *X_t_* ∈ ℝ^*n*^, for *t* = 1, …, *T*, each *A_k_* represents a coefficient matrix of dimension *n* × *n*, *ε_t_* is a zero mean Gaussian random vector of size *n* (white noise), and *p* denotes the maximal lag length. This model can also be rewritten in the form:

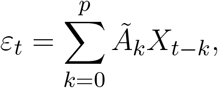

where *Ã*_0_ is an identity matrix and *Ã_k_* = −*A_k_* for *k* > 0. This can be transformed into the frequency domain:

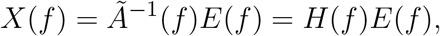

where *H*(*f*) is a non-symmetric transfer matrix of the system and *E*(*f*) is a noise component in a frequency domain. Then, Directed Transfer Function (see Kaminski and Blinowska (1991), Dauwels et al. (2010)) can be defined in terms of *H*:

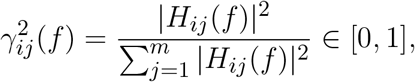

where the frequency-dependent normalization is chosen so that 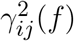 quantifies the fraction of (total) inflow to channel *i* that arrives from channel *j*. To obtain one value per link *j* → *i*, we averaged the 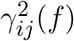 values through the frequencies *f* = 1, …, 256. DTF was computed in a sliding window of 100ms length with 50ms shift. Model order *p* for VAR(*p*) was put to 10 for all the patients, in order to decrease heterogeneity in the results.

### 5.4 Statistical analysis/detailed pipeline

The general diagram of the analysis pipeline is shown in Figure 4, detailed description of the pipeline is below. We denote:

- *T_DTF_* = {[−200:−100]ms, [−150:−50]ms, …, [700:800]ms} is a moving window for DTF computation; baseline and reaction subsets for DTF are *T_bsl_* = {[−200:−100]ms, [−150:−50]ms, …, [−50:50]ms} and *T_reac_* = {[0:100]ms, …, [700:800]ms},
- *T_PLV_* = [1, …, 510] is the time index for PLV; baseline and reaction subsets for PLV are *T_bsl_* = [1, …, 102] and *T_reac_* = [103, …, 512],
- 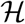 = [1:2]Hz, [2:4]Hz, [4:8]Hz, [8:16]Hz, [16:32]Hz, [32:64]Hz, [64:128]Hz is a set of frequency bands,
- *N^pat^* is a number of patients included in the analysis,
- *p* = 1, …, *N^pat^* = 15 is a patient number,
- *N*(*p*) is a number of analyzed channels for patient *p*,
- *i, j* are channel numbers, *i, j* = 1, …, *N*(*p*),
- *n*(*r*_1_, *r*_2_) is a number of connections between regions *r*_1_ and *r*_2_,
- *r*(.) ∈ {1, …, 7} is a function of channel number, providing the number of the ROI to which the channel belongs,
- *s* = *s*(*t, i*)[*p*] is the measured signal, set of multivariate time series for every patient *p*; *t* = 1, …, 512 is time index, and *i* is the index of the channel, *i* ∈ {1, …, *N* (*p*)}.
- *s_f_* = *s_f_* (*t, i*)[*p*] - measured signal *s*(*t, i*)[*p*], band-pass filtered to frequency band *f* ∈ 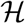.

For the measured data, we compute time-frequency resolved connectivity per patient between all channels *F* (*t, f*)[*i, j, p*] on the filtered signal *s_f_*

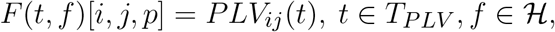

or

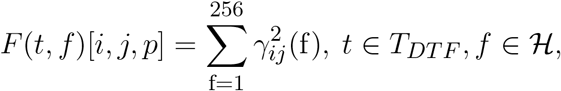

where *p* is a index of patient, *i, j*, *i* ≠ *j* are indices of channels *i, j* = 1, …, *N*(*p*) and *f* denotes the frequency band. Then, for signal of every subject *s*[*p*] we generate 100 multivariate Fourier surrogates from the baseline, i.e. the interval [−200:0] ms. Such surrogate data preserve the signal smoothness and correlation structure present in the (subject-, channel- and trial-specific) baseline. From the described surrogates, we computed the same time-frequency resolved connectivity *F_sim_*(*t, f, k*)[*i, j, p*] as we did for the data, *k* = 1, …, 100 is a surrogate index.

Based on the computed connectivity for data and surrogates, we created the time-frequency binary map of significance per pair of channels for data and corresponding surrogates *B*(*t, f*)[*i, j, p*] and *B_sim_*(*t, f*)[*i, j, p*]:

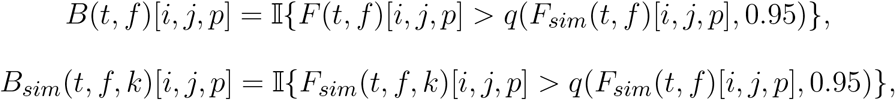

where 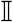 is an indicator function, *q*(*x, p*) is a *p*-level quantile of sample *x*.

Then, for every pair of ROIs, we created a connectivity density (*”heatmap”*) for data and surrogates by averaging all the binary maps for pair of ROIs:

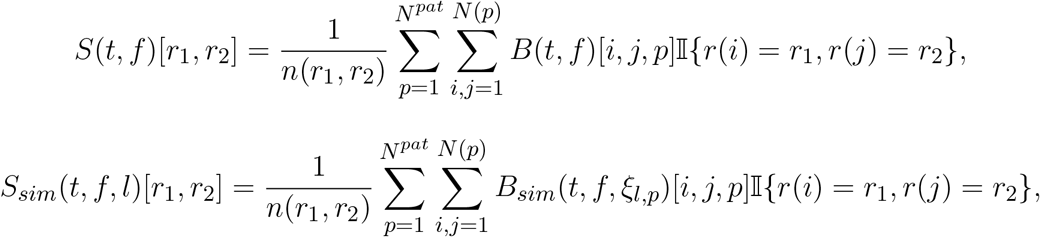

where 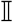 is an indicator function, *ξ_l,p_* is a random number indicating which of the 100 surrogate data for patient *p* should be used for the *l*-th simulated group heatmap, *r*(*i*) is number of ROI including channel *i*, *n*(*r*_1_, *r*_2_) is number of heatmaps for ROI pair *r*_1_ and *r*_2_, *t* ∈ *T_DTF_* or *t* ∈ *T_PLV_*, 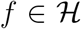, and *l* = 1, …, 1000 is number of simulated heatmaps.

Further, we computed mean reaction value *l*(*r*_1_, *r*_2_) as average heatmap in reaction window minus average heatmap in the baseline:

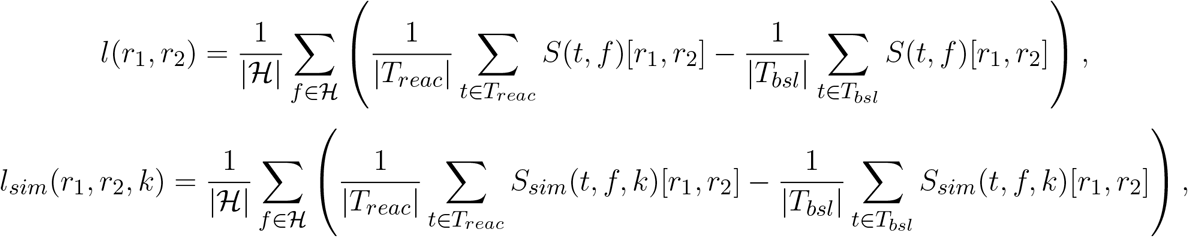

where *T_bsl_* and *T_reac_* are time points/windows of baseline and reaction, respectively, *r*_1_, *r*_2_ = 1, …, 7 are ROI indices, and *k* = 1, …, 1000 is number of simulations. Then, *p*-value for every pair of ROIs was computed by comparing the statistics *l* with the distribution *l_sim_*, obtained from surrogates as the percentile: *l* = *q*(*l_sim_, P* (*r*_1_, *r*_2_)). In the Results (Section 6) we discussed graphs, obtained from the matrix *P* by applying the Hochberg’s procedure (Hochberg, 1988) for controlling the family-wise error rate (FWE) at the 0.05 error level.

### 5.5 Software

The data preparation and the analysis were performed in MATLAB 2020b (Mathworks, Inc.). DTF was computed using MATLAB implementation (Omidvarnia, 2020; Omidvarnia et al., 2011). Vector autoregression coefficients were computed using the ARFIT package (Neumaier and Schneider, 2001). PLV was computed using MATLAB implementation Namburi (2011), based on Lachaux et al. (1999). Pictures of brain were generated via BrainNet (Xia et al., 2013).

## 6 RESULTS

We computed two general interaction graphs between the considered 7 ROIs: the undirected (based on PLV) and directed (DTF-based) connectivity network. The directed graph was obtained by DTF, the undirected one by PLV. Figure 5 shows the links between 7 ROIs with significant increase after the stimuli. The corresponding p-values can be found in Tables 2 and 3.

**Table 2.** P-values for testing the null hypothesis of no significant increase of the DTF during the reaction interval. Values less than 0.05 are marked in bold, significant values after the FWE-correction are marked with an asterisk (*). Matrix orientation is the following: from ROI in column to ROI in row. Thus, for example, the connection from ROI 6 to ROI 4 is significant uncorrected with the *p*-value 0.012.

| DTF | ROI 1 | ROI 2 | ROI 3 | ROI 4 | ROI 5 | ROI 6 | ROI 7 |
| --- | --- | --- | --- | --- | --- | --- | --- |
| ROI 1 | **0.006** | 0.054 | - | - | 0.128 | **0.029** | - |
| ROI 2 | 0.955 | 0.135 | 0.000 | - | - | 0.090 | 0.226 |
| ROI 3 | - | 0.141 | 0.706 | 0.090 | - | **0.000\*** | **0.042** |
| ROI 4 | - | - | 0.074 | 0.430 | - | **0.012** | 0.878 |
| ROI 5 | 0.841 | - | - | - | 0.119 | - | - |
| ROI 6 | **0.047** | 0.563 | 0.750 | 0.673 | - | **0.000\*** | - |
| ROI 7 | - | 0.937 | **0.015** | **0.006** | - | - | 0.900 |

**Table 3.** P-values for testing the null hypothesis of no significant increase of the PLV during the reaction interval. Values less than 0.05 are marked in bold, significant values after the FWE-correction are marked with an asterisk (*). We present only the upper triangle as the matrix is symmetric.

| PLV | ROI 1 | ROI 2 | ROI 3 | ROI 4 | ROI 5 | ROI 6 | ROI 7 |
| --- | --- | --- | --- | --- | --- | --- | --- |
| ROI 1 | **0.000\*** | **0.000\*** | - | - | **0.000\*** | **0.000\*** | - |
| ROI 2 | - | **0.000\*** | 0.565 | - | - | **0.000\*** | **0.011** |
| ROI 3 | - | - | **0.000\*** | **0.000\*** | - | **0.000\*** | **0.000\*** |
| ROI 4 | - | - | - | **0.001\*** | - | **0.000\*** | **0.001** |
| ROI 5 | - | - | - | - | **0.000\*** | - | - |
| ROI 6 | - | - | - | - | - | **0.000\*** | - |
| ROI 7 | - | - | - | - | - | - | **0.001\*** |

**Figure 5.**
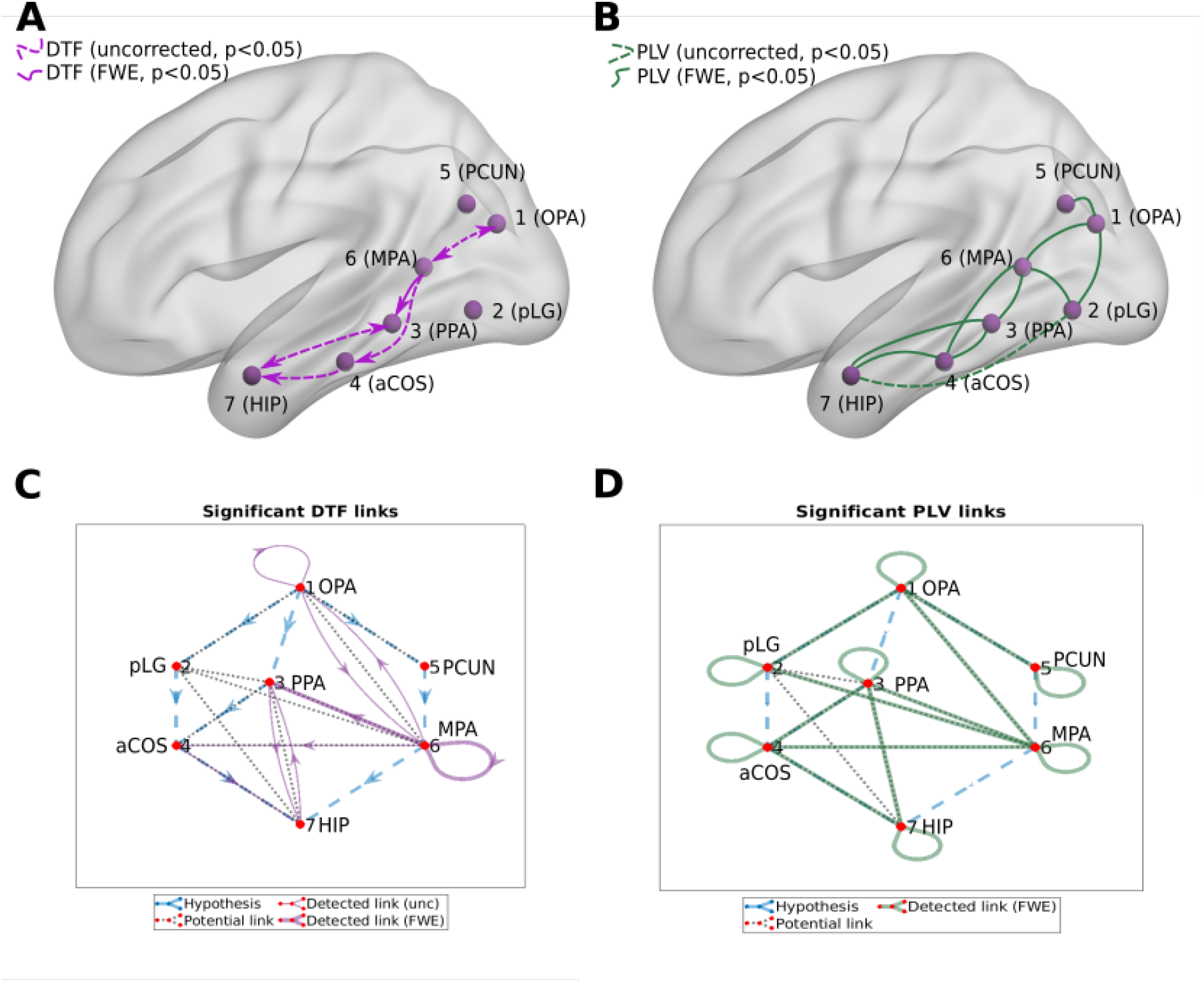
Connectivity graphs between the 7 ROIs, obtained by DTF (A and C panels, directed graphs, violet links) and PLV (B and D panels, undirected graphs, green links). Statistical significance was computed for the difference of the average heatmap value after and before the stimulus. Blue dashed lines in the C and D panels show the hypothesis of connections between ROI; black dotted lines show links presented at least in one subject; violet/green thin lines show significant links (uncorrected, *p* < 0.05); violet/green lines show significant links (FWE-corrected, *p* < 0.05).

Using PLV, we found a statistically significant increase in synchronization after the stimuli in all the assessed links between ROIs. In contrast, the causal inference created a much sparser network, with only one significant connection, when using the strict multiple comparison correction – The ROI 6 between the the dorsal and ventral streams influenced ROI 3, in the ventral stream. More connections were revealed without multiple comparison correction. In particular, in addition to the link from ROI 6 to ROI 3, we found increased connectivity in bidirectional interaction between ROIs 3 and 7, and between ROIs 1 and 6. In addition, the analysis revealed increase in unidirectional connectivity from ROI 6 to 4 and from ROI 4 to 7.

To reiterate, depending on the implantation scheme of each patient, the connectivity of each ROI with other ROIs was based on different channels from different patients. Figure 3 summarizes the number of patients and connections between channels in one or another ROI. To know specifically, from which brain regions the specific ROI connectivity emerged, we investigated in detail the localization of involved channels.

The connectivity from **ROI 6 to 3** was based on 9 connections between 10 channels. The six ROI 6 channels were localized mostly on the border between retrosplenial cortex (RSC) and either precuneus or the superior part of the lingual gyrus (sLG), while the four ROI 3 channels were all localized around the border between the anterior and posterior part of the parahippocampal place area (PPA), with the MNI y coordinate between −53 and −31 (−42 being the anterior/posterior boundary according to Baldassano et al. (2016)). The interaction between **ROIs 3 and 7** is based on data from 17 channels. Of them, the 12 channels in ROI 3 were primarily localized in the fusiform gyrus (FG) or lingual gyrus (LG), with MNI y coordinate mostly (10 of 12) posterior to −42, which again corresponds to posterior PPA. On the other hand, the 5 channels in ROI 7 were localized in the anterior hippocampus (HIP), entorhinal cortex (EC) and temporopolar gyrus, with MNI z coordinate below −19 (−9 being the anterior/posterior hippocampus boundary according to Morgan et al. (2011). The interaction between **ROIs 1 and 6** is based on data from one subject, with two channels in ROI 1, both localized between middle occipital gyrus and posterior angular gyrus (in transverse occipital sulcus), and four channels in ROI 6, all localized just between superior lingual gyrus (sLG) and retrosplenial cortex (RSC). The connectivity from **ROI 6 to 4** is based on data from two subjects, with four channels in ROI 6, all on the border of RSC and precuneus or LG, and four channels in ROI 4, all in posterior parahippocampal gyrus (PHG), with MNI y coordinate between −27 and −33 (the anterior/posterior border being at −20 according to Pruessner et al. (2002). The connectivity from **ROI 4 to 7** is based on data from 27 channels. Of them, 12 channels in ROI 4 were mainly localized in posterior part of PHG and adjacent FG, with MNI y coordinate posterior to −24, and 15 channels in ROI 7 were located primarily in the anterior HIP, EC and amygdala, with MNI z coordinate below −18.

Figures 6 and 7 present the heatmaps that were used for computation of graphs in Figure 5. We divided the heatmaps to two classes: connectivity from ROI to itself (Fig. 6), and connectivity between the different ROIs (Fig. 7). Recall that the heatmap of connectivity from ROI *i* to itself was estimated from the collection of connections between different channels, both of them laying in ROI *i*. Heatmap ROI *i* to ROI *j*, *i* ≠ *j* was computed from pairs of channels, for which the source channel belonged to ROI *i*, and the target channel was in ROI *j*.

**Figure 6.**
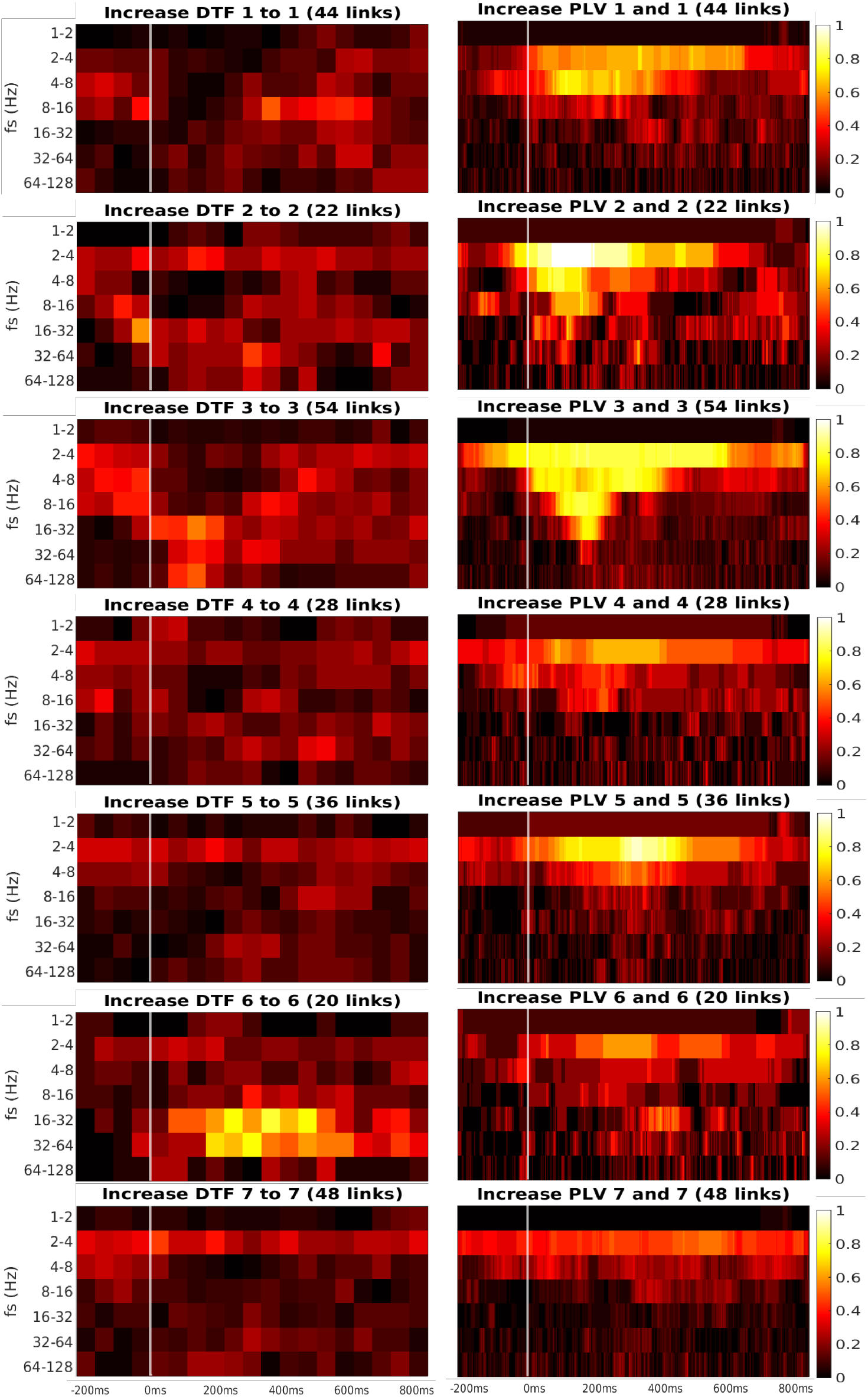
Time-frequency heatmaps of number of significant links per pair of ROI that are computed for the channels with significant reaction to the stimuli.

**Figure 7.**
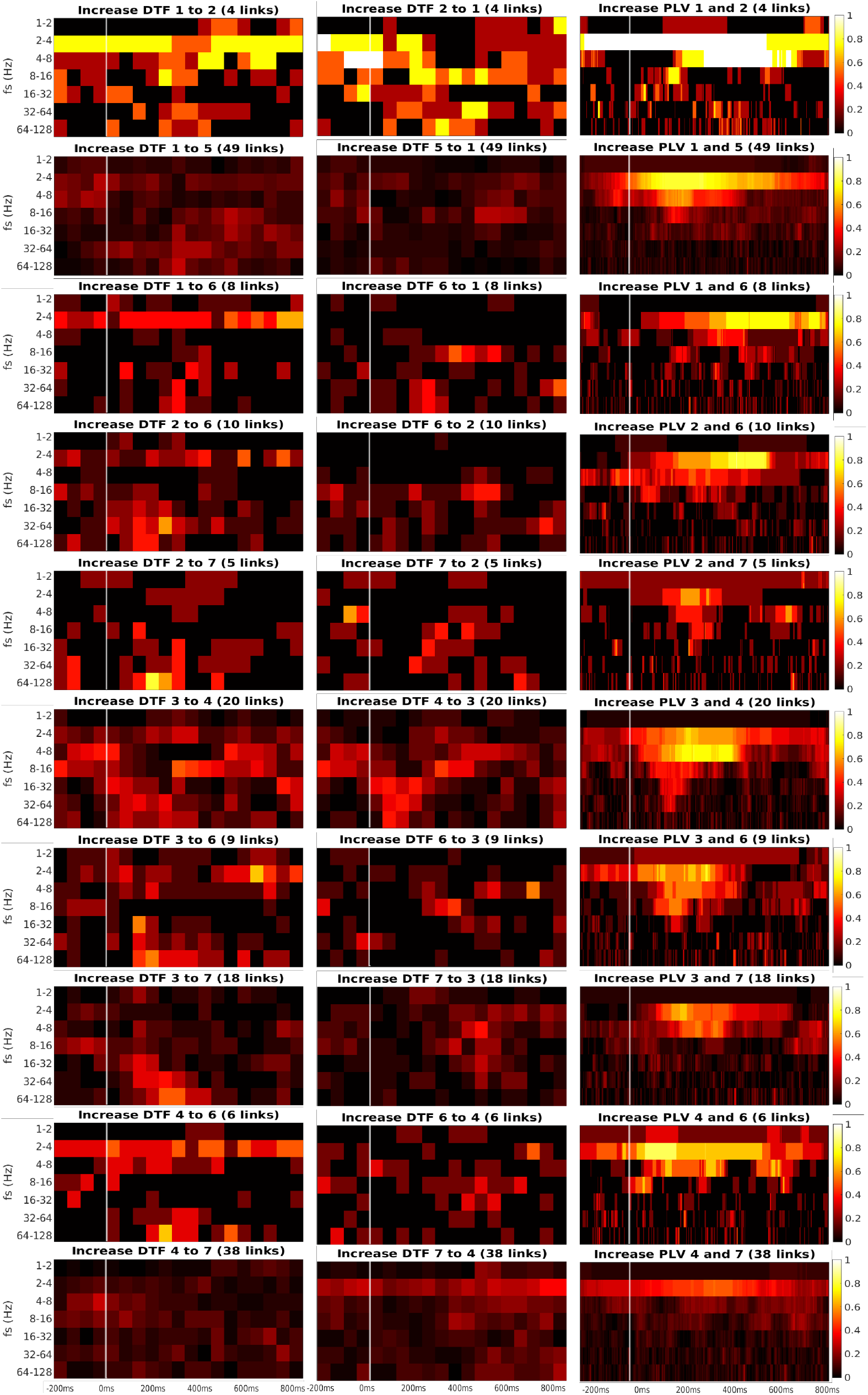
Time-frequency heatmaps of number of significant links per pair of ROI that are computed for the Channels with significant reaction to the stimuli.

Every heatmap shows the dynamical connectivity for several frequency bands (*y*-axis), with the time on the *x*-axis marking the start of sliding window for DTF, and the relevant time instant for PLV. Color of every entry corresponds to the percentage of significant links in particular time-frequency point between investigated pair of ROIs. Higher values of the heatmap (colour closer to yellow and white) mark a high proportion of links showing significantly increased connectivity and suggest at which time and frequency an increased communication emerges between the respective regions.

A robust significant increase in synchronization was often detected at the frequency bands 2 to 4 Hz and 4 to 8 Hz. The particular time interval of robust increase is difficult to define and seems to depend on the particular pair of ROIs. DTF increase is not as robust as in PLV, but generally, it also showed an increase at 2-4Hz frequency. DTF to/from ROI 3 and 6 seemed to have an increase at high frequencies, unlike the PLV, which showed a main increase at the low frequencies. Also, both of the connectivity measures showed a more robust increase inside the individual ROIs compared to the connectivity between ROIs. The presented heatmaps, however, are useful only for exploration purposes and should not be over-interpreted. The presented pictures element-wise were not associated with the statistical *p*-values. We also analyzed a connectivity decrease after the stimulation but didn’t find significant results.

## 7 DISCUSSION

As stated in the introduction, the inference of brain graph analysis from iEEG data faces an extensive set of challenges. We have outlined these in the introduction, proceeding from those that are shared across a large set of inference situations, to those that are relatively specific to the situation of iEEG - namely the partial, sparse and heterogeneous sampling of the relevant brain network in any single patient. Based on the discussion of the possible ways to tackle these challenges, we have made a range of methodological decisions in order to build up a pipeline allowing robust statistical inference of the alterations of brain connectivity network during processing visual stimuli. The application of the pipeline was demonstrated on an example dataset, giving rise to an group-level network in terms of both functional and effective connectivity.

Note that the functional and effective connectivity correspond to different yet related concepts - the true underlying interactions (in principle captured by the effective connectivity) can be considered as the the cause of the synchronization of the brain activity. However the true interactions that are to be estimated by effective connectivity methods might not necessarily happen exactly between the pair of regions for which synchrony is observed. Indeed, due to common drivers or mediators, two regions can be synchronized in their activity, while not interacting directly.

Also, while we have made a relatively common choice of connectivity measures from each family, the results could surely be (potentially substantially) different for other choices of connectivity measures. In particular, the functional connectivity measure used (i.e., PLV), is in two important aspects different from the classical counterpart of DTF in the functional connectivity family, that is, from the coherence. In particular, the PLV works with phase only (unlike the DTF and coherence, which depend on both the phase and amplitude of the signal). Secondly, the PLV measures the synchronization across trials, rather than synchronization across time (as would mean phase coherence do). We have had two-fold motivation for this choice of functional connectivity selection: firstly, PLV is an widespread and commonly used method. Secondly, the heterogeneity of connectivity methods used helps to illustrate the wide applicability of the pipeline. Of course, the framework is in principle applicable along with a range of other connectivity methods, including the natural alternative to DTF, i.e. the partial directed coherence Baccalá et al. (2001); Baccalá and Sameshima (2021); Sameshima and Baccalá (1999). Yet, one may surely argue for the application of many alternative methods from the plethora of the existing ones.

Although the specific application was not the main rationale of this study, the utilization of the developed pipeline provided interesting results. In the studied case of network changes in reaction to the visual stimuli, the developed methodology allowed us to robustly detect several interesting phenomena.

First of all, we confirmed the involvement of the parieto-medial temporal pathway in scene perception, translating visuospatial information between dorsal and ventral visual streams during visual scene analysis. This pathway connects the parietal lobe with locations in the medial temporal lobe (ROI 3 and 4), via MPA (ROI 6, also called retrosplenial complex). The existence of this pathway was suggested based on rat studies showing that associative parietal cortex lesions affect the firing of place cells in the medial temporal lobe (Save et al., 2005). The pathway is claimed to include also the retrosplenial region (Byrne et al., 2007). Based on anatomical data, it connects the angular gyrus (homological to caudal inferior parietal lobule in animals) in the parietal lobe to the medial temporal lobe either directly or indirectly, with the indirect connection leading via the retrosplenial cortex and posterior cingulate cortex (Kravitz et al., 2011). Both direct and indirection connections were also demonstrated in humans using resting-state fMRI data (Boccia et al., 2017). Our data show that this indirect pathway (from ROI 6 to ROIs 3 and 4) is active also during static visual scenes processing in humans. In agreement with another connectivity study (Baldassano et al., 2016), we show the MPA connection with the PPA (ROI 3), around the border between its anterior and posterior parts.

Second, we confirmed the anterior hippocampal region (ROI 7) connectivity with more posterior areas in the medial temporal lobe: the parahippocampal place area (PPA, ROI 3) and region around the anterior collateral sulcus (ROI 4). The hippocampus (HIP) is a medial temporal structure associated primarily with declarative and spatial memory, but its function and connectivity vary along its long anterior-posterior axis (Strange et al., 2014). While the function of the posterior portion (pHIP) seems to be related to spatial memory, the anterior part (aHIP) has been associated with emotional processing, novelty detection, and semantic memory. Accordingly, there are connectivity differences, with the aHIP being linked, among others, to the amygdala, ventral tegmental area and hypothalamus, while pHIP is coupled to posterior parietal areas and retrosplenial cortex (Poppenk et al., 2013). However, there seems to be some overlap, as the aHIP shows connection also to more posterior areas, like anterior PPA and MPA (Boccia et al., 2017), in one study even larger than pHIP (Baldassano et al., 2016). Extending these studies, our current results document a connection of aHIP (ROI 7) with even posterior part of PPA (ROI 3). The hippocampal connections to the parahippocampal gyrus (PHG) also differ between its anterior to posterior part. While aHIP shows preferential connectivity with the perirhinal cortex (anterior part of PHG), pHIP is preferentially connected to the parahippocampal cortex (posterior part of PHG) (Libby et al., 2012). However, this preferential connectivity pattern does not exclude the functional coupling of other areas. Our data show information flow from posterior PHG (ROI 4) to aHIP (ROI 7). However, our collection of channels did not contain pHIP. Therefore, for both posterior PPA and parahippocampal cortex, we could not exclude their indirect association with aHIP via pHIP.

Third, we found the reciprocal information flow between ROI 1 in OPA and ROI 6 in MPA. At the same time, we hypothesized to find connectivity between ROI 1 and PPA (ROI 3), but we found no patients with both ROIs implanted in our dataset. Related to that, a recent resting-state fMRI connectivity study (Baldassano et al., 2016) distinguished two scene processing networks. The anterior one connects angular gyrus with retrosplenial cortex, anterior portion of PPA and anterior HIP. In contrast, the posterior one consists of OPA and posterior portion of PPA. In agreement with this study, the two channels in our ROI 1, showing connectivity with ROI 6 in our data, seem to be a part of the anterior scene processing network, which is related more to memory and navigation. In fact, these two channels in ROI 1 were localized still in medial occipital gyrus, but deep at the anterior end of transverse occipital sulcus, thus on the posterior margin of angular gyrus. Therefore, our data support the separation of anterior and posterior scene processing networks (Baldassano et al., 2016).

Albeit the methodology was tailored to the nature of the analyzed data, the results achieved are still clearly limited by some aspects of the data. For instance, the low spatial coverage, in particular the lack of implantation of posterior occipital cortex, prevented us from tracking the early development of the visual signal processing. Low number of patients involved not only affected the statistical power of the inference, but indirectly also the amount of pairs of regions that we were able to test for interactions, and together with the relative sparseness of the implantation it generally decreased the spatial resolution of the estimated network. Indeed, the regions were arguably large enough so that they would be functionally heterogeneous. On the other side, should one decide to work with smaller regions, there might not be sufficient number of subjects with implantation spanning any given pair of regions, leaving thus each connection inference specific to a single subject.

Needless to say, the results in principle might be affected by any residual differences in the cognitive processing, and, more importantly, brain dynamics in the epilepsy patients compared to healthy subjects. Although the volunteering patients all had normal vision, and any segments with detected interictal epileptiform activity (or seizures) were removed from the analyzed data, one can never exclude the possibility of some systematic bias in neuronal dynamics and functional neuroanatomy due to the chronic brain disease. Indeed, the selection of the regions here was based on a data driven approach used in a previously published study; an alternative approach used in the field is to apply some standard anatomical atlas.

## AUTHOR CONTRIBUTIONS

AP, KV and JHl conceptualized and validated the study. Data curation: AP, AK, JHa, KV and JHl. Formal analysis was performed by AP and JHl. Funding acquisition: PM, KV and JHl. Methodology: AP, JHa, PS, KV and JHl. Project administration: AP, PM, KV and JHl. Resources: PM, AK, KV and JHl. Software and visualisation: AP. Supervision: JHl. AP, KV, JHl were writing the original draft; all the authors contributed to the investigation and writing - review and editing - the manuscript.

## ETHICS STATEMENT

The studies involving human participants were reviewed and approved by Ethics Committee of Motol University Hospital (02-APR-2014). Written informed consent to participate in this study was provided by the participant or participants’ legal guardian/next of kin..

## ACKNOWLEDGMENT

We thank all patients for their participation, and Nad’a Bednárová, Helena Buchtová, Lucie Paterová, Iveta Fajnerová, Tereza Nekovářová and Jana Kalinová for their help with collecting the iEEG data.

## DATA AVAILABILITY STATEMENT

Pipeline scripts (in MATLAB) and the estimated connectivity matrices are available upon request from the corresponding author.

